# Unexpected diversity of CPR bacteria and nanoarchaea in the rare biosphere of rhizosphere-associated grassland soil

**DOI:** 10.1101/2020.07.13.194282

**Authors:** Alexa M. Nicolas, Alexander L. Jaffe, Erin E. Nuccio, Michiko E. Taga, Mary K. Firestone, Jillian F. Banfield

**Affiliations:** Department of Plant and Microbial Biology, University of California, Berkeley, Berkeley, CA; Nuclear and Chemical Sciences Division, Lawrence Livermore National Laboratory, Livermore, CA, USA; Department of Environmental Science, Policy, and Management, University of California, Berkeley, Berkeley, CA; Earth and Environmental Sciences, Lawrence Berkeley National Laboratory, Berkeley, CA; Department of Earth and Planetary Science, University of California, Berkeley, Berkeley, CA; Chan Zuckerberg Biohub, San Francisco, CA; Innovative Genomics Institute, University of California, Berkeley, Berkeley, CA

## Abstract

Candidate Phyla Radiation (CPR) bacteria and nanoarchaea populate most ecosystems, but are rarely detected in soil. We concentrated particles less than 0.2 *μ*m from grassland soil, enabling targeted metagenomic analysis of these organisms, which are almost totally unexplored in soil. We recovered a diversity of CPR bacteria and some nanoarchaea sequences, but no sequences from other cellular organisms. The sampled sequences include Doudnabacteria (SM2F11) and Pacearchaeota, organisms not previously reported in soil, as well as Saccharibacteria, Parcubacteria and Microgenomates. CPR and DPANN (an acronym of the names of the first included archaea phyla) enrichments of 100-1000-fold were achieved compared to bulk soil, in which we estimate these organisms comprise about 1 to 100 cells per gram of soil. Like most CPR and DPANN sequenced to date, we predict these microorganisms live symbiotic, anaerobic lifestyles. However, Saccharibacteria, Parcubacteria, and Doudnabacteria genomes sampled here also encode ubiquinol oxidase operons that may have been acquired from other bacteria, likely during adaptation to aerobic soil environments. We posit that although present at low abundance, CPR bacteria and DPANN archaea could impact overall soil microbial community function by modulating host organism abundances and activity.

## Introduction

Interactions among soil microorganisms impact biogeochemical cycling and overall ecosystem function. Recent metagenomic analysis of soil microbial communities revealed that many steps of key reaction pathways central to transformations in soil are partitioned among coexisting organisms (1). Other interactions are mediated by molecules such as vitamins and antimicrobial compounds (2–4). Further, there is the potential for a variety of symbiotic interactions, including those that involve obligate reliance on coexisting organisms for even the most basic requirements (5,6). Candidate Phyla Radiation (CPR) bacteria and DPANN archaea (an acronym of the names of the first included phyla: Diapherotrites, Parvarchaeota, Aenigmarchaeota, Nanoarchaeota and Nanohaloarchaeota) are detected across ecosystems and are often predicted to be obligate, anaerobic (epi)symbionts (5,7) that depend on other organisms for basic cellular building blocks (8). However, CPR bacteria and DPANN archaea have rarely been identified in soil (1,9,10). This may be because isolation-based methods fail for organisms unable to grow alone and primers used in 16S rRNA gene surveys can have mismatches that preclude detection (11). Genome-resolved metagenomic analyses circumvent these limitations, yet there are few reports of CPR metagenome-assembled genomes (MAGs) from soil (12), almost certainly because of the low relative abundance of these bacteria. Here, we tested the hypothesis that CPR and DPANN have been overlooked in the soil rare biosphere by taking advantage of the expected very small cell sizes of these organisms (13). Using methods developed to isolate viruses from soil, we investigated the soil nanoparticulate fraction for nanobacteria and nanoarchaea. Thus, we sequenced concentrated-soil-effluent that had passed through a 0.2 *μ*m filter used to remove larger cells (14,15), and recovered a diversity of nanobacterial and nanoarchaeal sequences.

## Results and Discussion

We sampled rhizosphere-associated soil from the top 10 cm of an annual grassland from the UC Research and Extension Center near Hopland, California in February 2018. We added buffer and collected the effluent from a subset of the soil samples, which we passed through a 0.2 *μ*m-filter, concentrated, and treated with DNase to remove extracellular DNA that could have derived from larger lysed cells (**Fig 1a**, **Supplementary Methods)**. To evaluate enrichment, bulk DNA was extracted from the same soil samples for whole community shotgun DNA sequencing, generating what are here referred to as “bulk metagenomes.” Approximately 20 Gbp of sequence was obtained from each of six concentrates and two bulk samples. In addition to recovering viral sequences and mobile elements from these small-particle-concentrate metagenomes, we reconstructed sequences from CPR and nanoarchaea genomes. From these data, we resolved 26 draft genomes that were > 70% complete (estimated using a CPR-specific single copy gene set (11)), with < 10% contamination derived from either CPR or DPANN. No CPR or DPANN genomes were assembled from the bulk metagenomes.

**Figure 1.**
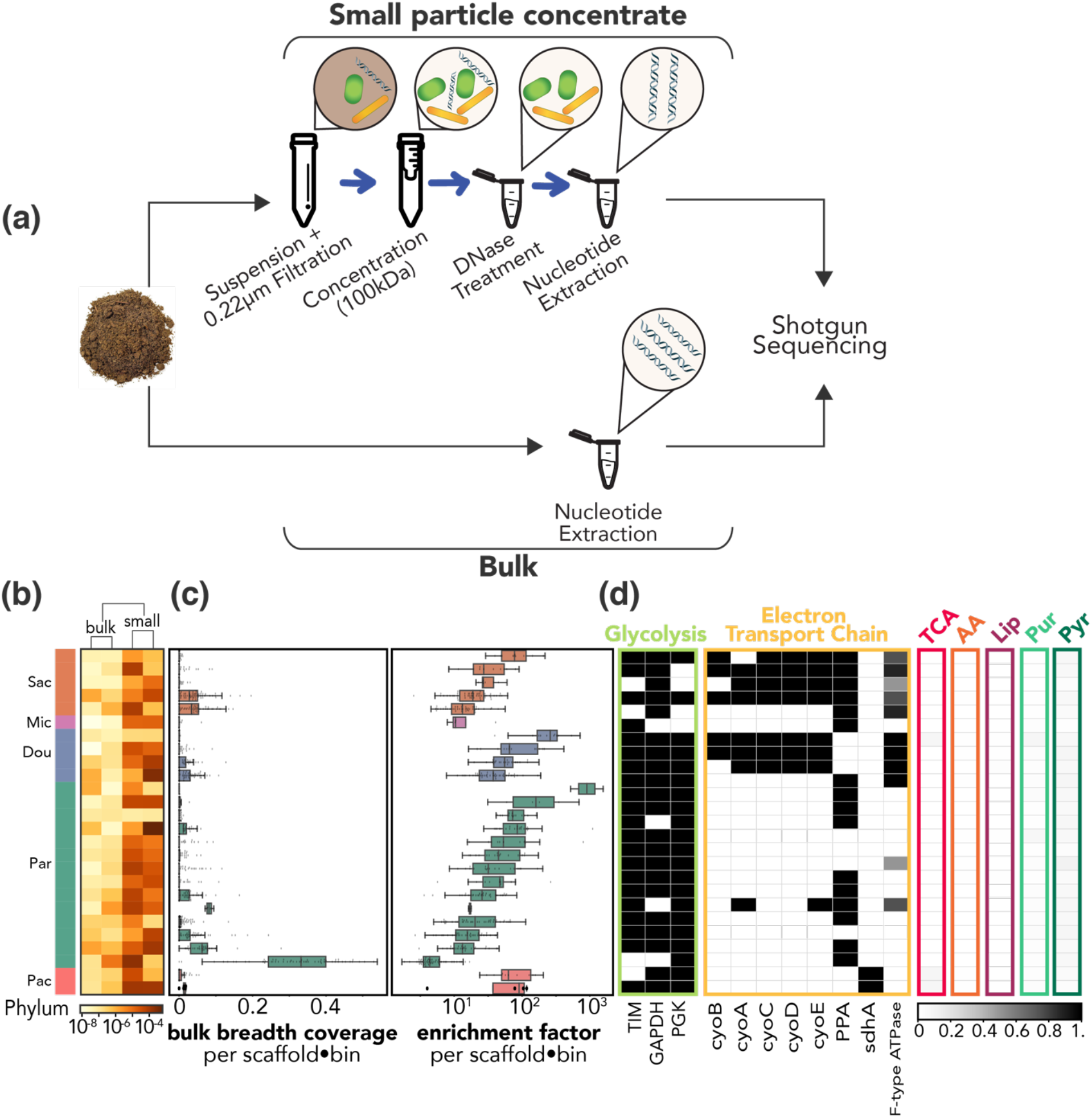
Enrichment and metabolic profiles of CPR in soil concentrate metagenomes. **(a)** Method for concentration of small particles from soil for metagenomic sequencing (*top*) compared to sample preparation methods for bulk soil metagenomes (*bottom*). **(b)** Heatmap showing relative abundance of 26 organisms by phylum (Sac: Saccharibacteria, Mic: Microgenomates, Dou: Doudnabacteria, Par: Parcubacteria, Pac: Pacearchaeota) across bulk metagenomes and concentrate metagenomes. **(c)** Coverage based metrics showing recovery and enrichment in all concentrates combined, relative to bulk fractions combined. *Left*: Breadth of coverage of scaffolds comprising each genome (bin) in the bulk fraction. *Right:* Enrichment factor (i.e., relative abundance of a scaffold from the concentrate metagenome over a scaffold’s bulk metagenome relative abundance) for each genome. **(d)** Metabolic analysis of each genome including: presence of each of three glycolysis genes that are highly conserved among CPR; genes involved in electron transport chain, including percentage completeness of F-type ATPase; and percentage completeness (grey scale) of the TCA cycle (tricarboxylic acid cycle) and pathways for AA (amino acid biosynthesis), Lip (lipid biosynthesis), Pur (purine biosynthesis), Pyr (pyrimidine biosynthesis).

We found that sequences from cells < 0.2 *μ*m were almost exclusively from 15 lineages of CPR bacteria and one DPANN archaea phylum (**Fig 2**; **Supplementary Fig. 1**). Importantly, these bacteria and archaea were only detected in bulk metagenomes from the same soil samples by read mapping to small particle concentrate-derived genomes. In fact, most sequences that comprise each CPR or DPANN genome were completely absent in bulk metagenome samples, and bulk metagenome reads mapping to concentrate-derived CPR genomes provided little to no genome coverage of the assembled sequences (**Fig. 1c)**. Further, from the 74 bacterial 16S rRNA sequences recovered from the concentrate metagenomes, all of which were assigned to CPR lineages, we predict more than half (42 16S rRNA gene sequences) would not have been detected using standard amplicon sequencing primers. Notable was the unexpected phylum-level diversity of CPR lineages in the concentrate metagenomes. Previously, a genome of TM7 (Saccharibacteria) was reported from the same soil, but sampled at less than 1x coverage from bulk soil (6), and Microgenomates and Parcubacteria have been genomically sampled twice at low abundance (12,16). To our knowledge, this is the first report of Pacearchaeota, Doudnabacteria and potentially a novel clade of Saccharibacteria in soil.

**Figure 2.**
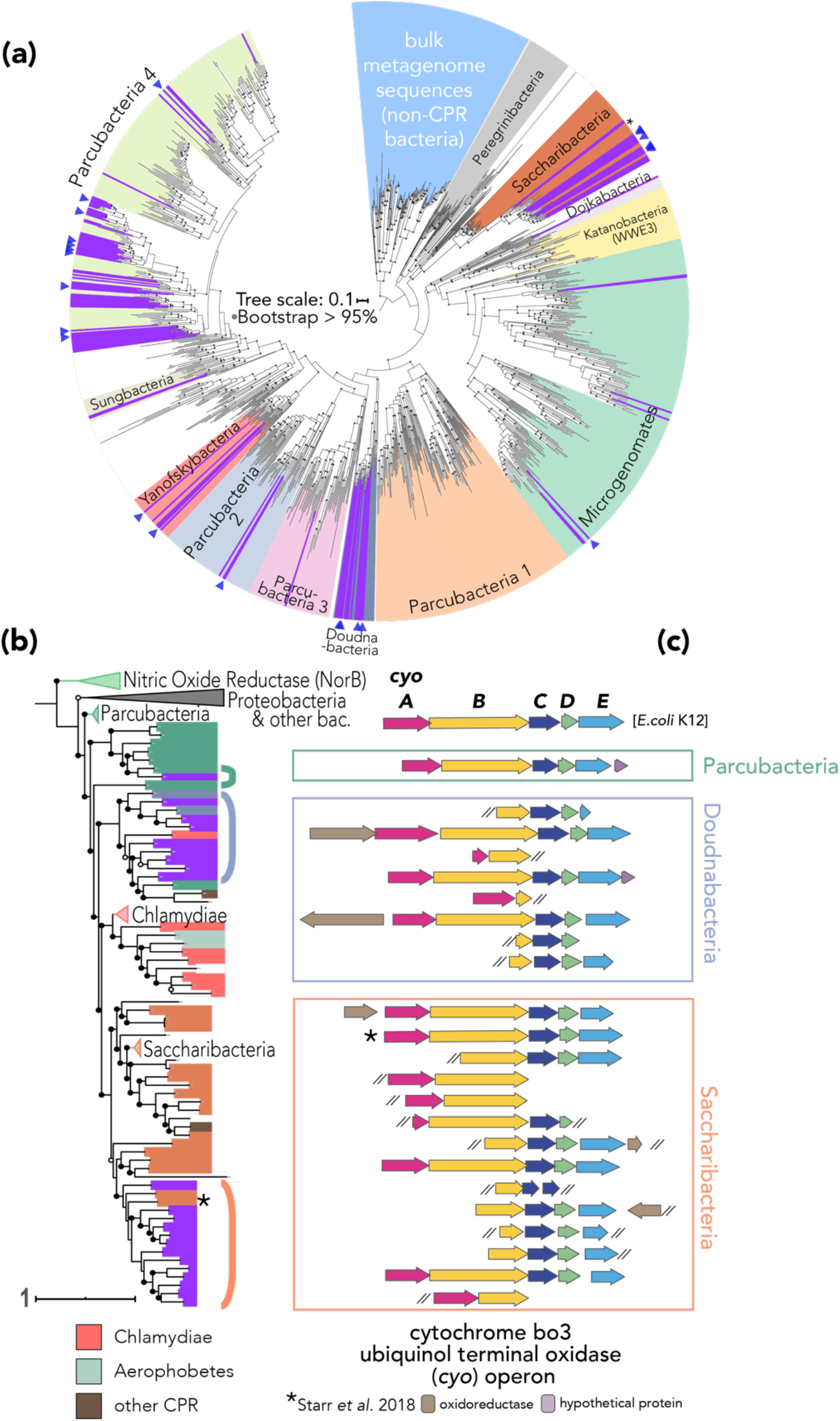
Soil CPR phylogeny and cytochrome operon synteny. Sequences assembled from the small concentrate metagenomes are shown in purple. **(a)** *rpS3* tree of CPR bacteria rooted using *rpS3* sequences that were assembled from bulk metagenomes, in light blue. Blue triangles denote draft genome recovered. Nodes with bootstrap values greater than or equal to 0.95 are marked as filled gray circles. **(b)** Phylogenetic relationships of cytochrome bo3 ubiquinol terminal oxidase subunit I across bacterial phyla. Circles overlaid on nodes correspond to support values (unfilled >0.50, filled >0.70). The asterisked sequences show the placement of *T. rhizospherense cyoB*, and its operon in (c) (5). Brackets next to tree tips correspond to phyla by color (green: Parcubacteria, blue: Doudnabacteria, orange: Saccharibacteria) and to sequence order in (c) synteny diagram. Tree rooted using a heme-copper oxidase superfamily member, the nitric oxide reductase (*norB*). **(c)** Synteny diagram of cytochrome ubiquinol oxidase operon genes (*cyoA, cyoB, cyoC, cyoD,* and *cyoE*) with operon from *E. coli* K12 as reference. Scale bars correspond to the average number of substitutions per site across alignment.

Comparing sequence coverage from concentrates to that from the bulk metagenomes, we calculate that filtration enriched the relative abundance of genomes by 100-1000x (**Fig 1c)**. We approximate, given the relative abundance of the most and least abundant CPR genomes in each of the bulk and concentrate metagenome, that CPR cells may comprise on the order of 1-100 cells per gram of soil. Given estimates of 10^9^ microbial cells per gram of soil, this would equate to, at maximum, ~10^−5^ percent of microbial cells in a gram of soil (**Supplementary Methods**).

Given the unique challenges of the soil environment for microbes, we next assessed whether these soil CPR and DPANN exhibited similar traits to their counterparts in other environments. Recent studies show that CPR bacteria generally appear to have the capacity for glycolysis and fermentation (17), but often lack complete pathways to predictably *de novo* synthesize nucleotides and have many gaps in metabolism that suggest an obligate symbiotic lifestyle (8). We find that most of the genomes from this sampling effort encode the three central glycolysis enzymes reportedly found in nearly all CPR bacteria: triose phosphate isomerase (TIM), glyceraldehyde 3-phosphate (GAPDH), and phosphoglycerate kinase (PGK) (17). As expected, based on prior studies of CPR bacteria and DPANN archaea (8), the genomes contain few if any TCA cycle genes and lack genes of the electron transport chain, and for synthesis of lipids and nucleotides **(Fig 1d)**, suggesting they live anaerobic lifestyles with strong dependence on resources from other organisms. However, we identified an operon encoding a multi-subunit cytochrome bo3 ubiquinol terminal oxidase in genomes of three Doudnabacteria, eight Saccharibacteria, and one Parcubacteria as well as in non-binned CPR phyla sequences from the concentrate metagenomes. Previously, Starr *et al.* (2018) found an operon similar to those reported here in the first Saccharibacteria genome described from soil, *T. rhizospherense* (**Fig 2c**). A few CPR loci also included an ORF annotated as an oxidoreductase or a conserved hypothetical protein. It remains unclear whether the combination of this ubiquinol oxidase and the associated oxidoreductase (**Fig 2c**) confers the ability to use O_2_ in some way. While, to our knowledge, this is the first report of this operon in Doudnabacteria, finding it in these lineages indicates that some form of aerobic respiratory capacity may be common in soil-associated Saccharibacteria specifically, and perhaps soil CPR more broadly. Our synteny analysis **(Fig 2c)** shows identical gene order for the cyo operon as in the highly studied *E. coli* K12 operon (18) and in the *T. rhizospherense* genome (6), although some CPR loci were incomplete due to assembly fragmentation. Due to the potential of these genes to encode for metabolic processes that detoxify reactive oxygen species, we hypothesize that alternatively this operon may confer an adaptive advantage for CPR to live in aerophilic environments such as surface soil. We generated a maximum-likelihood tree of subunit 1 (*cyoB*) of the *cyo* operon to test whether the operon exhibited a pattern of vertical inheritance in our CPR genomes (**supplementary methods**; **Fig 2b)**. This phylogenetic analysis suggests that this gene cluster has been laterally transferred from other bacteria such as Proteobacteria or Chlamydiae into these CPR at least once, with perhaps different origins for gene clusters from Parcubacteria and Doudnabacteria compared to those from Saccharibacteria. Further, based on *cyoB* phylogeny, *T. rhizospherense cyoB* appears more closely related to Saccharibacteria sequences from this study compared to *rpS3* phylogeny, which may further underscore local adaptation to soil.

## Conclusion

Here, we conducted a targeted study of CPR and nanoarchaea in a soil ecosystem to expand our understanding of soil-dwelling microbes. Using typical sequencing allocations for soil metagenomics, we were only able to recover genomes for these little-known community members through concentrating small particles from soil. The results suggest that CPR bacteria and nanoarchaea are relatively rare, but present in soil. We detected multi-subunit cytochrome operons in three CPR lineages, possibly conferring adaptation to aerophilic surface soil environments. While the precise ecological roles of these organisms in situ remains unclear, their predicted requirement for interaction with nearby community members to satisfy their metabolic and nutritional needs, and their previously reported close physical association with other cells (5,7) suggest that they may play a still undescribed role in soil microbial interaction networks. Further, the capacity to selectively filter for genomes suggests that if these organisms attach to larger microbial cells, the association can be physically disrupted. We suggest that like other small, low abundant community members such as bacteriophages and parasites of eukaryotes, CPR and nanoarchaea likely contribute to nutrient recycling in soil and impact the functioning of associated bacteria and thus affect communities at large.

## Acknowledgements

This research was supported by the U.S. Department of Energy Office of Science, Office of Biological and Environmental Research Genomic Science program under Awards DE-SC0020163 and DE-SC0010570 to MKF and the NIH grant DP2AI117984 to MET. Work conducted at Lawrence Livermore National Laboratory was supported by the U.S. Department of Energy Office of Science, Office of Biological and Environmental Research Genomic Science program under Award SCW1678, LLNL Lab Directed Research and Development Award 18-ERD-041, and under the auspices of the U.S. DOE under Contract DE-AC52-07NA27344. We thank Cindy J. Castelle for support inferring archaea phylogenetic relationships.

## References

1. Diamond S, Andeer PF, Li Z, Crits-Christoph A, Burstein D, Anantharaman K, et al. Mediterranean grassland soil C–N compound turnover is dependent on rainfall and depth, and is mediated by genomically divergent microorganisms. Nat Microbiol [Internet]. 2019 Aug 20 [cited 2019 Dec 4];4(8):1356–67. Available from: https://doi.org/10.1038/s41564-019-0449-y

2. Jiang X, Zerfaß C, Feng S, Eichmann R, Asally M, Schäfer P, et al. Impact of spatial organization on a novel auxotrophic interaction among soil microbes. ISME J [Internet]. 2018 Jun 23 [cited 2018 May 17];12(6):1443–56. Available from: http://www.nature.com/articles/s41396-018-0095-z

3. Abrudan MI, Smakman F, Grimbergen AJ, Westhoff S, Miller EL, van Wezel GP, et al. Socially mediated induction and suppression of antibiosis during bacterial coexistence. Proc Natl Acad Sci U S A [Internet]. 2015;112(35):1504076112–. Available from: http://www.pnas.org/content/early/2015/07/22/1504076112.short?rss=1

4. Abreu NA, Taga ME. Decoding molecular interactions in microbial communities. FEMS Microbiol Rev. 2016;019(40):648–63.

5. He X, McLean JS, Edlund A, Yooseph S, Hall AP, Liu S-Y, et al. Cultivation of a human-associated TM7 phylotype reveals a reduced genome and epibiotic parasitic lifestyle. Proc Natl Acad Sci [Internet]. 2015 Jan 6 [cited 2020 Jan 27];112(1):244–9. Available from: http://www.pnas.org/lookup/doi/10.1073/pnas.1419038112

6. Starr EP, Shi S, Blazewicz SJ, Probst AJ, Herman DJ, Firestone MK, et al. Stable isotope informed genome-resolved metagenomics reveals that Saccharibacteria utilize microbially-processed plant-derived carbon. Microbiome [Internet]. 2018 Dec 3 [cited 2018 Jul 3];6(1):122. Available from: https://microbiomejournal.biomedcentral.com/articles/10.1186/s40168-018-0499-z

7. He CY, Keren R, Whittaker M, Farag IF, Doudna JA, Cate JH, et al. Huge and variable diversity of episymbiotic CPR bacteria and DPANN archaea in groundwater ecosystems. bioRxiv. 2020;2020.05.14.094862.

8. Castelle CJ, Brown CT, Anantharaman K, Probst AJ, Huang RH, Banfield JF. Biosynthetic capacity, metabolic variety and unusual biology in the CPR and DPANN radiations. Nat Rev Microbiol [Internet]. 2018 Oct 4 [cited 2020 Jan 26];16(10):629–45. Available from: http://www.nature.com/articles/s41579-018-0076-2

9. Portillo MC, Leff JW, Lauber CL, Fierer N. Cell size distributions of soil bacterial and archaeal taxa. Appl Environ Microbiol [Internet]. 2013 Dec 15 [cited 2018 May 9];79(24):7610–7. Available from: http://fiererlab.org/wp-content/uploads/2014/09/Portillo_etal_AEM_2013.pdf

10. Delgado-Baquerizo M, Oliverio AM, Brewer TE, Benavent-González A, Eldridge DJ, Bardgett RD, et al. A global atlas of the dominant bacteria found in soil. Science (80-) [Internet]. 2018 Jan 19 [cited 2020 Jan 14];359(6373):320–5. Available from: https://www.sciencemag.org/lookup/doi/10.1126/science.aap9516

11. Brown CT, Hug LA, Thomas BC, Sharon I, Castelle CJ, Singh A, et al. Unusual biology across a group comprising more than 15% of domain Bacteria. Nature [Internet]. 2015 Jul 15 [cited 2018 May 6];523(7559):208–11. Available from: http://www.nature.com/articles/nature14486

12. Kroeger ME, Delmont TO, Eren AM, Meyer KM, Guo J, Khan K, et al. New biological insights into how deforestation in amazonia affects soil microbial communities using metagenomics and metagenome-assembled genomes. Front Microbiol. 2018 Jul 23;9(JUL):1635.

13. Luef B, Frischkorn KR, Wrighton KC, Holman H-YN, Birarda G, Thomas BC, et al. Diverse uncultivated ultra-small bacterial cells in groundwater. Nat Commun [Internet]. 2015 May 27 [cited 2019 Jul 30];6(1):6372. Available from: http://www.nature.com/articles/ncomms7372

14. Trubl G, Solonenko N, Chittick L, Solonenko SA, Rich VI, Sullivan MB. Optimization of viral resuspension methods for carbon-rich soils along a permafrost thaw gradient. 2016;(C):1–24.

15. Williamson KE, Corzo KA, Drissi CL, Buckingham JM, Thompson CP, Helton RR. Estimates of viral abundance in soils are strongly influenced by extraction and enumeration methods. Biol Fertil Soils. 2013;49(7):857–69.

16. Sharrar AM, Crits-Christoph A, Méheust R, Diamond S, Starr EP, Banfield JF. Bacterial secondary metabolite biosynthetic potential in soil varies with phylum, depth, and vegetation type. BioRxiv. 2019;

17. Jaffe AL, Castelle CJ, Matheus Carnevali PB, Gribaldo S, Banfield JF. The rise of diversity in metabolic platforms across the Candidate Phyla Radiation. BMC Biol [Internet]. 2020 Dec 19;18(1):69. Available from: https://www.biorxiv.org/content/10.1101/2019.12.18.881540v1

18. Abramson J, Riistama S, Larsson G, Jasaitis A, Svensson-Ek M, Laakkonen L, et al. The structure of the ubiquinol oxidase from Escherichia coli and its ubiquinone binding site. Nat Struct Biol. 2000;7(10):910–7.

