## Supplementart Materials & Methods for "Unexpected diversity of CPR bacteria and nanoarchaea in the rare biosphere of rhizosphere-associated grassland soil"

### SUPPLEMENTARY METHODS

#### Soil Sampling and DNA extraction

Soil samples were taken on February 16, 2018 from the top 10 cm of six field plots grown with *Avena barbata* as the dominant vegetation as part of an ongoing field experiment at the University of California Hopland Research and Extension Center. From two of these plots, 0.4 g of root- and stone-free soil were separated for DNA extraction as described in Nuccio *et al.* 2016 (1). For all six samples, nanoparticulate concentrates were produced starting with 10 g of soil as in Trubl *et al.* 2016 (2), described in brief with modifications here. An "amended" potassium citrate buffer (1% potassium citrate, 1x PBS, 100 mM $MgSO_4$, pH 7) was added to the soil 1 mL per g soil (to supersaturate soil) in a 50 mL falcon tube. The mixture was homogenized by shaking manually and placed on a horizontal shaker at 400 rpm for 15 minutes. Tubes were loaded onto a horizontal tube holder and vortexed for 3 minutes. Following vortexing, tubes were manually homogenized for 30 seconds and then centrifuged for 10 minutes at 4,700 g in a swinging bucket rotor at 4 ˚C to pellet soil. After pipetting supernatants off into fresh tubes, buffer was added to soil pellets in the same ratio previously described and steps were repeated twice more, each time adding the supernatant to the same collecting tube. In total, buffer was added to each soil sample three times. The aggregated supernatant in the collection tube was filtered using a Steriflip, which contains a 0.22 $\mu$m PES membrane vacuum filter and stored overnight at 4 ˚C.

Next, the nearly 30 mL of filtered supernatant was concentrated for DNase treatment and subsequent DNA extraction using an Amicon Ultra-15 Centrifugal Filter Unit with a 100 kDa molecular weight cutoff. First, Amicon filters were blocked using 2 mL of 1% BSA (0.2 $\mu$m filter-sterilized) to coat the filter and were incubated for 1 hour at 4˚C. Following incubation, filters were centrifuged in a swinging bucket rotor at 1,500 g for 10 minutes (or until all BSA passes through the filter). Excess BSA was then removed both from the reservoir and filter. Filters were then washed with 2 mL 1x PBS and similarly centrifuged to remove PBS. Samples were added to coated and washed Amicon devices and centrifuged 5 to 10 minutes at a time until samples were concentrated to approximately 250 $\mu$L. Concentrated samples were removed from filter reservoirs by adding 250 $\mu$L of amended potassium citrate buffer in three steps (amounting to a total nanoparticulate concentrate of 1 mL). After 250 $\mu$L of buffer was added to a reservoir, the Amicon device was vortexed for 30 seconds and then the concentrate in the reservoir was pipetted up and down, washing the walls of the membrane and the concentrate was added to a fresh 2 mL Eppendorf tube, and the step was repeated until the Eppendorf tube contained 1 mL of filtered and concentrated small particles from soil. DNase treatment was carried out as in Hurwitz *et al.* 2013 (3).

Following DNase treatment to rid nanoparticulate concentrates of free DNA, we extracted DNA from small cells and particles after an iron flocculation step, as described in John *et al.* 2011 (4), combined with a phenol:chloroform strategy based on that of Bergallo *et al.* 2006 (5). In short,

1 μL of 0.02 μm filter-sterilized Iron Chloride Solution (0.1 g Fe per 1 mL deionized water) was added per mL of each sample. The mixture was centrifuged at 14,000 g for 20 minutes and the supernatant was discarded, with care to avoid disturbing the pellet. The pellet was resuspended in 20 μL of 0.02 μm filter-sterilized Ascorbic-EDTA buffer (0.2 M EDTA, 0.4 M ascorbic acid, pH 6-7) and mixed by pipetting and vortexing until the pellet was disaggregated.

To extract DNA from the 0.2 μm size-filtered cell and particulate fraction of soil, 250 μL of Phenol:Chloroform:Isoamyl Alcohol (25:24:1) (pH 8) was added to each sample and vortexed for 1 minute. Samples were then incubated on ice for 15 minutes, vortexing occasionally to homogenize samples, and centrifuged at 14,000 g for 5 minutes. The top layer of the aqueous phase was transferred to a Phase Lock Gel tube and 250 μL of chloroform was added to this tube. Tubes were centrifuged at 14,000 g for 5 minutes and supernatant was transferred to a fresh Eppendorf tube. 25 μL of 3 M sodium acetate (pH 5) and 1.5 μL of glycoblue were added to the supernatant and mixed. Next, 250 μL of isopropanol was added and mixed thoroughly and samples were incubated at -80 ˚C for 20 minutes. Following incubation, samples were centrifuged at 14,000 g for 20 minutes and supernatant was discarded. The pellet was washed with 500 μL of freshly prepared 70% ethanol and centrifuged at 14,000 g for 5 minutes. Supernatant was removed from each sample, and samples were centrifuged for an additional 1 minute at 14,000 g. Using a fine tipped pipet, remaining ethanol was removed from the pellet, and samples were dried for 5 minutes. Finally, the pellet was resuspended in 15 μL of Tris-HCl. DNA was quantified from all samples by Qubit dsDNA High Sensitivity Assay Kit.

**DNA Sequencing, assembly, and genome reconstruction**

All metagenomic library preparation and DNA sequencing was performed at the Vincent J. Coates Genomics Sequencing Laboratory at UC Berkeley. Library preparation was carried out using Accel-NGS 1S Plus kit (Swift Biosciences) for sequencing on an Illumina HiSeq4000 platform producing 150 bp paired-end reads. Raw sequencing data were processed with the Illumina supported CASAVA bcl2fastq (v2.19) program.

Following removal of adapter sequences and other sequencing contaminants using BBTools, quality trimming with Sickle, and quality control with FastQC, sequences were assembled using IDBA-UD. Specifically, 2 co-assemblies were generated: read files of all nanoparticulate fractionated samples were concatenated for coassembly to establish the small particle concentrate metagenome assembly and read files for all bulk metagenomes were concatenated to create the bulk metagenome assembly. Open reading frames (ORFs) for each scaffold were predicted using Prodigal, and functional predictions per ORF were made through similarity searches using USEARCH to query KEGG, UniRef100, and UniProt databases using default parameters. Full metagenome samples and their annotations were then uploaded into our in-house analysis platform, ggKbase (https://ggkbase.berkeley.edu). High-quality draft bacterial and archaeal genomes were chosen by DASTool (6) from bins created by differential coverage binners CONCOCT (7), MaxBin (8), MetaBAT (9), and manually using the ggKbase interface based on GC content, DNA sequence coverage, and taxonomic affiliation. Scaffold taxonomic affiliation represents the consensus taxonomy (>50%) of annotations of all predicted ORFs on a scaffold. Following DASTool selection of bins, genomes were further manually curated on ggKbase and checked for completeness and contamination by running CheckM (10) with a custom set of single copy genes for CPR genomes (11). Archaeal genome completeness was predicted based on 38 single copy archaeal genes (12). Only the 26 genomes predicted to be greater than 70% complete and with less than 10% contamination by single copy genes are reported. These genomes are available at: https://ggkbase.berkeley.edu/soilcpr.

**Relative abundance of draft genomes, breadth coverage, and enrichment ratios**

To calculate relative abundance of each draft CPR and archaeal genome, reads from each of the six nanoparticulate fraction metagenomes and the two bulk metagenomes were mapped to the set of 26 draft genomes using Bowtie2. Mapped reads were then filtered using CoverM (https://github.com/wwood/CoverM) to ensure all reads mapped shared at least 97% minimum identity with genome sequences. Relative abundance per genome, as visualized using the seaborn package in Python to generate a heatmap, was calculated by aggregating all read counts mapping to scaffolds that comprised bins for the bulk metagenomes and their paired small particle concentrate metagenomes and normalizing by the total number of reads for each metagenome.

To determine how the small particle concentrate metagenomes better recover CPR and nanoarchaea metagenomes we established two metrics: (1) bulk breadth coverage (2) enrichment factor. Bulk breadth coverage was calculated by mapping all concatenated bulk metagenome reads to CPR and nanoarchaea genome sequences using Bowtie2, followed by filtering reads with a minimum identity threshold of 97% and calculating the breadth of a sequence reads mapped to i.e. “covered fraction” using CoverM. The distribution of breadth coverage for each scaffold in a bin was visualized using seaborn boxplots laid over swarmplots. Second, the enrichment factor was calculated by computing an overall relative abundance for each scaffold of a draft genome from all small particle concentrate reads divided by the overall relative abundance for each scaffold from bulk metagenome reads, and was similarly visualized using seaborn.

**CPR and DPANN approximation of rare biosphere**

In order to estimate the percent of soil microbial cells comprised of CPR or DPANN, we compared the amount of DNA extracted from the bulk and small particle concentrate metagenomes to the most and least relative abundant genomes. Calculation details and assumptions are shown in **Supplementary Table 1**.

Estimated count of CPR or nanoarchaea cells in soil

| Bulk metagenome | Small particle concentrate metagenome |
|---|---|
| **Total DNA extracted** | **Total DNA extracted** |
| ~3 µg/g soil = $3\times10^{-6}$ g/g soil | ~100 ng/g soil = $1\times10^{-7}$ g / g soil |
| **Relative abundance of most abundant CPR** | **Relative abundance of most abundant CPR** |
| $10^{-6}$ | $10^{-4}$ |
| **Mass of DNA/Mbp** | |
| $10^{-15}$g | |
| **~Mass of 1 Cell** | |
| ~$5\times10^{-15}$g | |
| **Max # cells per g soil** | **Max # cells per g soil** |
| $\frac{(10^{-6} \bullet 3\times10^{-6} g)}{5\times10^{-15} g}$ = 600 cells | $\frac{(10^{-4} \bullet 1\times10^{-7} g)}{5\times10^{-15} g}$ = 2000 cells |
| **Least abundant lineage** | **Least abundant lineage** |
| $10^{-8}$ | $10^{-7}$ |
| **Min # cells per g soil** | **Min # cells per g soil** |
| $\frac{(10^{-8} \bullet 3\times10^{-6} g)}{5\times10^{-15} g}$ = 6 cells | $\frac{(10^{-7} \bullet 1\times10^{-7} g)}{5\times10^{-15} g}$ = 2 cells |

**Supplementary Table 1 I Estimated count of CPR or nanoarchaea cells in soil.** *Left* shows approximation of the number of CPR or DPANN cells of a given lineage in a gram of soil using bulk metagenome parameters. *Right* provides the same estimate using parameters derived from the small particle concentrate metagenome.

### Metabolic overview of draft genomes

In order to assess the metabolic capacity of draft genomes reported here, KoFamScan (13) was run to assign KEGG orthology (KO) numbers to draft genome ORFs. For glycolysis, we subsequently searched ORF annotations for three genes present in nearly all CPR genomes (14) (triose phosphate isomerase (TIM): K01803, glyceraldehyde 3-phosphate (GAPDH): K00134, and phosphoglycerate kinase (PGK): K00927). To define the tricarboxylic acid (TCA) cycle by presence of enzymes we used a set of KOs as described in Méheust et al. 2019 (15). To query for presence of electron transport enzymes in these draft genomes, we searched for all KO numbers that comprise the oxidative phosphorylation pathway, map00190. Since all oxidative phosphorylation enzymes found were either part of the F-type ATPase module (m00157) or were single enzymes present from a module, we grouped the F-type ATPase module in the heatmap to visualize its completion, and other enzymes present were included in the heatmap by their presence/absence. To determine whether these genomes maintained the capacity to synthesize nucleotides *de novo*, we looked for the presence of enzymes of the pyrimidine (map00240) and purine biosynthesis pathways (map00230). Given the sparseness of the results, we collapsed the presence of purine (pur) and pyrimidine (pyr) enzymes into their respective categories out of the total number of genes included in each of the pathways. For amino acid (AA) biosynthesis we searched for enzymes that were part of alanine, aspartate and glutamate metabolism (map00250); glycine, serine and threonine metabolism (map00260); cysteine and methionine metabolism (map00270); valine, leucine and isoleucine biosynthesis (map00290); lysine biosynthesis (map00300); arginine biosynthesis (map00220); arginine and proline metabolism (map00330); histidine metabolism (map00340); tyrosine metabolism (map00350); phenylalanine metabolism (map00360); tryptophan metabolism (map00380); and phenylalanine, tyrosine and tryptophan biosynthesis (map00400). Similarly, these separate pathways were combined into the single category of AA biosynthesis by calculating how many of the total unique enzymes were present in each genome. Lipid (Lip) biosynthesis was calculated in the same way, combining the sparse results of queried enzymes of the following lipid biosynthesis pathways: glycerolipid metabolism (map00561), glycerophospholipid metabolism (map00564), and biosynthesis of unsaturated fatty acids (map01040). A heatmap was then generated using the Seaborn Python package to show gene presence/absence or percent pathway completeness.

**Cytochrome bo3 ubiquinol terminal oxidase (*cyo*) operon curation and phylogenetic analysis**

Cytochrome bo3 ubiquinol terminal oxidase (*cyo*) operon was identified through KOfam matches to the cytochrome o ubiquinol oxidase module (M00417) and homology searches using ggKbase. Scaffolds containing ubiquinol oxidase subunits chosen for further analysis had a consensus taxonomy of Bacteria, and contained subunit 1 (*cyoB*, K02298, EC:7.1.1.3) plus an additional subunit of the ubiquinol oxidase directly upstream or downstream (22 sequences in total). To construct the synteny diagram, each of these selected scaffolds was then searched for additional ORFs considered part of the operon. These ORFs were identified by being on the same sense strand and directly adjacent to or overlapping with, i.e. without gaps, an ORF identified as part of the operon. ORFs were included despite the presence of a gap if the annotated ORF had homology to ORFs that were directly adjacent to *cyo*-annotated ORFs in the context of other scaffolds.

For the phylogenetic analysis of the 22 cytochrome ubiquinol oxidase subunit I sequences, we created a reference set of sequences from: (1) BLAST hits (max target seqs set to 100 and a minimum evalue of 1E-100) which were then clustered using USEARCH to remove duplicate hits (0.99 identity); (2) Searching 175 evenly sampled bacteria lineages (16) using the K02298 KOfam with a threshold score of 700; and for additional CPR sequences (3) K02298 queried against a phylogenetically comprehensive database of CPR genomes (16) with the same threshold. Additionally, nitric oxide reductase subunit B (NorB) Swiss-prot sequences were downloaded from UniProt (April 22, 2020) and clustered to 0.80 identity using USEARCH to reduce redundancy among sequences. All sequences were aligned using MAFFT and trimmed using the mask feature in GeneiousPrime to strip columns with more than 95% gaps (729 amino acids). A maximum likelihood tree was then inferred for this alignment using IQTREE with ultrafast bootstrapping (-m TEST -nt AUTO -st AA -bb 1500) (16). To assign names to tree leaves, the Entrez module from the Biopython was used to query NCBI accession numbers to return sequence names and phylum-level lineage identification when possible. This identifier information was added to tree leaves using the ETE Toolkit (17). The resulting tree (and all trees described here) was visualized in iTOL (18).

**Phylogenetic classification of sequences**

To establish phylogeny of prokaryote sequences and genomes assembled from the small particle concentrate as compared to sequences recovered in the bulk metagenome, we used the S3 ribosomal protein (rpS3). rpS3 sequences were identified by an HMM search using Pfam (PF00189) from both bulk and small concentrate metagenome assemblies using the gathering threshold as the score cutoff. For the bacterial tree, we established a set of PF00189 hits from the set of CPR genomes curated in Jaffe *et al.* 2020 (14). An alignment of rpS3 sequences from known CPR (14), bulk metagenome, and small particle concentrates was generated using MAFFT, and trimmed using GeneiousPrime to strip columns with greater than 95% gaps (286 amino acids). The alignment was used to create a maximum likelihood tree inferred using IQTREE with ultrafast bootstrapping as previously described. The tree was rooted with rpS3 sequences derived from the bulk metagenome.

Finding that several rpS3 sequences from the small particle fraction were likely of archaeal origin, we aligned these sequences with rpS3 sequences from previously reported archaeal genomes (19). This alignment was trimmed initially using GeneiousPrime stripping columns with more than 95% gaps, and then manually (196 amino acids) and an approximately-maximum-likelihood phylogenetic tree was generated using FastTree. Reference sequences were then refined to better recapitulate previously found archaea phylogenetic relationships by subsetting reference archaeal sequences to three DPANN phyla: Diapherotrites, Pacearcheota, and Woesearcheota. This reduced alignment was used to infer a maximum likelihood tree using IQTREE with ultrafast bootstrapping with the same parameters previously described.

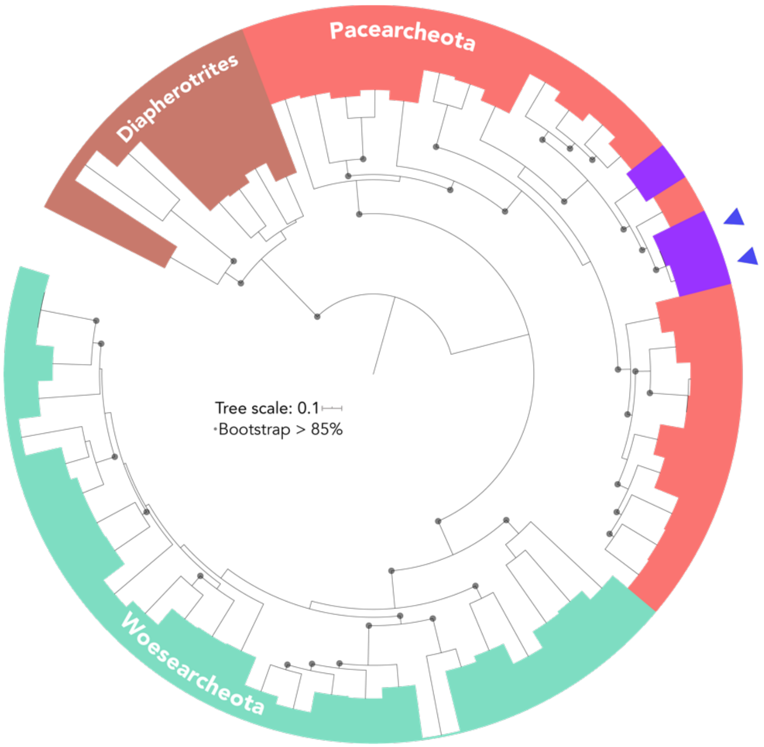


**Supplementary Figure 1 | Soil nanoarchaea phylogeny.** Maximum-likelihood tree of *rpS3* sequences of nanoarchaea. Purple colored sequences were assembled from the small concentrate metagenomes reported. Tree was rooted using *rpS3* sequences of Diapherotrite phylum sequences. Blue triangles represent *rpS3* sequences for which draft genomes were recovered. Nodes with bootstrap values, >85%, are marked as filled circles. Tree scale bar corresponds to the average number of substitutions per site across alignment.

**Supplementary References**


1. Nuccio EE, Anderson-Furgeson J, Estera KY, Pett-Ridge J, De Valpine P, Brodie EL, et al. Climate and edaphic controllers influence rhizosphere community assembly for a wild annual grass. Ecology. 2016;97(5):1307–18.
2. Trubl G, Solonenko N, Chittick L, Solonenko SA, Rich VI, Sullivan MB. Optimization of viral resuspension methods for carbon-rich soils along a permafrost thaw gradient. 2016;(C):1–24.
3. Hurwitz BL, Deng L, Poulos BT, Sullivan MB. Evaluation of methods to concentrate and purify ocean virus communities through comparative, replicated metagenomics. Environ Microbiol [Internet]. 2013 May [cited 2017 Feb 16];15(5):1428–40. Available from: http://doi.wiley.com/10.1111/j.1462-2920.2012.02836.x
4. John SG, Mendez CB, Deng L, Poulos B, Kauffman AKM, Kern S, et al. A simple and efficient method for concentration of ocean viruses by chemical flocculation. Environ Microbiol Rep [Internet]. 2011 Apr 1 [cited 2020 Apr 3];3(2):195–202. Available from: http://doi.wiley.com/10.1111/j.1758-2229.2010.00208.x
5. Bergallo M, Costa C, Gribaudo G, Tarallo S, Baro S, Ponzi AN, et al. Evaluation of six methods for extraction and purification of viral DNA from urine and serum samples. New Microbiol [Internet]. 2006 [cited 2018 Jan 16];29(2):111–9. Available from: http://www.newmicrobiologica.org/pub/allegati_pdf/2006/2/micro2_05_bergallo.pdf
6. Sieber CMK, Probst AJ, Sharrar A, Thomas BC, Hess M, Tringe SG, et al. Recovery of genomes from metagenomes via a dereplication, aggregation and scoring strategy. Nat Microbiol [Internet]. 2018 Jul 28 [cited 2019 May 12];3(7):836–43. Available from: http://www.nature.com/articles/s41564-018-0171-1
7. Alneberg J, Bjarnason BS, De Bruijn I, Schirmer M, Quick J, Ijaz UZ, et al. Binning metagenomic contigs by coverage and composition. Nat Methods [Internet]. 2014 Oct 30 [cited 2020 Jul 2];11(11):1144–6. Available from: https://github.com/
8. Wu Y-W, Tang Y-H, Tringe SG, Simmons BA, Singer SW. MaxBin: an automated binning method to recover individual genomes from metagenomes using an expectation-maximization algorithm. Microbiome [Internet]. 2014 Dec 1 [cited 2019 Mar 15];2(1):26. Available from: https://microbiomejournal.biomedcentral.com/articles/10.1186/2049-2618-2-26
9. Kang DD, Froula J, Egan R, Wang Z. MetaBAT, an efficient tool for accurately reconstructing single genomes from complex microbial communities. PeerJ [Internet]. 2015 Aug 27 [cited 2020 Jul 2];3(8):e1165. Available from: https://bitbucket.org/
10. Parks DH, Imelfort M, Skennerton CT, Hugenholtz P, Tyson GW. CheckM: assessing the quality of microbial genomes recovered from isolates, single cells, and metagenomes. Genome Res [Internet]. 2015 Jul [cited 2018 May 7];25(7):1043–55. Available from: http://www.ncbi.nlm.nih.gov/pubmed/25977477
11. Brown CT, Hug LA, Thomas BC, Sharon I, Castelle CJ, Singh A, et al. Unusual biology across a group comprising more than 15% of domain Bacteria. Nature [Internet]. 2015 Jul 15 [cited 2018 May 6];523(7559):208–11. Available from: http://www.nature.com/articles/nature14486
12. Probst AJ, Ladd B, Jarett JK, Geller-Mcgrath DE, Sieber CMK, Emerson JB, et al.

Differential depth distribution of microbial function and putative symbionts through sediment-hosted aquifers in the deep terrestrial subsurface. Nat Microbiol [Internet]. 2018 Mar 29 [cited 2020 Feb 2];3(3):328–36. Available from: http://www.nature.com/articles/s41564-017-0098-y
13. Aramaki T, Blanc-Mathieu R, Endo H, Ohkubo K, Kanehisa M, Goto S, et al. KofamKOALA: KEGG Ortholog assignment based on profile HMM and adaptive score threshold. Valencia A, editor. Bioinformatics [Internet]. 2020 Apr 1 [cited 2020 Jul 2];36(7):2251–2. Available from: https://academic.oup.com/bioinformatics/article-abstract/36/7/2251/5631907
14. Jaffe AL, Castelle CJ, Matheus Carnevali PB, Gribaldo S, Banfield JF. The rise of diversity in metabolic platforms across the Candidate Phyla Radiation. BMC Biol [Internet]. 2020 Dec 19;18(1):69. Available from: https://www.biorxiv.org/content/10.1101/2019.12.18.881540v1
15. Méheust R, Castelle CJ, Carnevali PBM, Farag IF, He C, Chen L-X, et al. Aquatic Elusimicrobia are metabolically diverse compared to gut microbiome Elusimicrobia and some have novel nitrogenase-like gene clusters. bioRxiv. 2019;765248.
16. Nguyen L-T, Schmidt HA, Von Haeseler A, Minh BQ. IQ-TREE: A Fast and Effective Stochastic Algorithm for Estimating Maximum-Likelihood Phylogenies. 2014 [cited 2020 Jul 2]; Available from: www.mbe.oxfordjournals.org/
17. Huerta-Cepas J, Serra F, Bork P. ETE 3: Reconstruction, Analysis, and Visualization of Phylogenomic Data. [cited 2020 Jul 2]; Available from: http://creativecommons.
18. Letunic I, Bork P. Interactive Tree Of Life (iTOL) v4: recent updates and new developments. Web Serv issue Publ online [Internet]. 2019 [cited 2020 Jul 2];47. Available from: https://itol.embl.de/version
19. Castelle CJ, Brown CT, Anantharaman K, Probst AJ, Huang RH, Banfield JF. Biosynthetic capacity, metabolic variety and unusual biology in the CPR and DPANN radiations. Nat Rev Microbiol [Internet]. 2018 Oct 4 [cited 2020 Jan 26];16(10):629–45. Available from: http://www.nature.com/articles/s41579-018-0076-2
